# Developmental differences in canonical cortical networks: insights from microstructure-informed tractography

**DOI:** 10.1101/2023.10.30.564863

**Authors:** Sila Genc, Simona Schiavi, Maxime Chamberland, Chantal Tax, Erika Raven, Alessandro Daducci, Derek K Jones

## Abstract

There is a growing interest in incorporating white matter fibre-specific microstructural properties into structural connectomes to obtain a more quantitative assessment of brain connectivity. In a developmental sample aged 8-18 years, we studied age-related patterns of microstructure-informed network properties locally and globally. First, we computed the diffusion-weighted signal fraction associated with each tractography-reconstructed streamline. Then, we generated microstructure-informed connectomes from diffusion MRI data using the convex optimization modelling for microstructure-informed tractography (COMMIT) approach. Finally, we estimated network characteristics in eight functionally defined networks (visual, somatomotor, dorsal attention, ventral attention, limbic, frontoparietal, default mode and subcortical networks). Our findings reveal that throughout child and adolescent development, global efficiency increases in the visual, somatomotor, and default mode networks, and mean strength increases in the somatomotor and visual networks. Nodes belonging to the dorsal and ventral visual pathways demonstrate the largest age-dependence in local efficiency, supporting previous evidence of protracted maturation of dorsal and ventral visual pathways. Our results provide compelling evidence that there is a prolonged development of visual association cortices.

## 1. Introduction

The transition from childhood to adolescence is a period of profound neurobiological and cognitive development where the human brain undergoes significant changes to refine neural substrates prior to adulthood (Blakemore & Choudhury, 2006). Essential to this process are the white matter pathways that form a structural scaffold facilitating connections and communication between cortical regions. Their development follows a stereotypical pattern of myelination, which closely mirrors the functional capacity of neural systems. For example, primary sensory, motor and visual pathways typically complete myelination by the first two years of life (Deoni et al., 2015), whereas frontal and temporal association regions continue to develop well into adulthood, with peak myelination happening in the second decade of life (Bartzokis et al., 2012; Yakovlev & Lecours, 1967). The process of axonal development is less clear, with early *ex vivo* studies indicating stabilization of corpus callosum axonal count by six months of age (LaMantia & Rakic, 1990) and further work indicating changes to axonal and myelin properties at pubertal onset (Genc et al., 2023; Juraska & Willing, 2017; Paus, 2010).

Developmental studies using magnetic resonance imaging (MRI) have revealed that white matter volume steadily increases over childhood and adolescence (Giedd et al., 1999; Lenroot & Giedd, 2006), likely by way of coupled radial growth of the axon and myelin sheath. In tandem, functional MRI (fMRI) studies suggest a greater degree of temporal network connectivity, which remodels from infancy to early adulthood (Grayson & Fair, 2017). Early in childhood, sensorimotor systems become well integrated and coordinated, and show little change into adulthood (Gu et al., 2015). Later in adolescence, functional hubs such as fronto-parietal, attentional and salience networks become increasingly segregated, allowing for flexibility as the adolescent brain becomes more adaptable to increase performance and efficiency (Bassett et al., 2011).

Diffusion magnetic resonance imaging (dMRI) has enabled novel discoveries in spatial and temporal patterns of white matter fibre development (Geeraert et al., 2019; Genc et al., 2018; Herting et al., 2017; Lebel & Beaulieu, 2011; Palmer et al., 2022; Tamnes et al., 2018). Structural connectivity has been studied using diffusion MRI tractography (Hagmann et al., 2007) to reconstruct white matter pathways or connections between nodes of interest (e.g., between distinct predefined cortical regions). Connection strength is commonly defined using white matter streamline count, i.e., the number of streamlines, derived from tractography, that run between nodes. However, this notion can be arbitrary, since streamline count is not biologically informative and can heavily depend on acquisition and processing parameters (Jones et al., 2013; Yeh et al., 2021; Zhang et al., 2022). Recent studies have attempted to improve the *status quo* in determining biologically informative determinants of connection strength using diffusion MRI (Smith et al., 2020; Zhang et al., 2022), however, the question remains: which measures are optimally informative?

To define more informative edge weights for the structural connectome, the ‘tractometry’ approach was introduced in (Bells et al., 2011; Jones et al., 2006; Kanaan et al., 2006) and employed to study typical white matter development (Chamberland et al., 2019). This approach includes the mapping of microstructural measures along tractography-reconstructed pathways and computing average values for quantitative comparisons between measures. A challenge arises when multiple bundles pass through the same imaging voxel (an extremely prevalent phenomena; see Jeurissen et al. (2013); Schilling et al. (2022)) which leads to biased measures assigned to each constituent bundle (Schiavi et al., 2022). The Convex Optimization Modelling for Microstructure Informed Tractography (COMMIT) (Daducci et al., 2015; Daducci et al., 2013) approach address this problem by deconvolving specific microstructural features on each streamline to recover individual contributions to the measured signal. By replacing the commonly used streamline count with intra-axonal signal fraction (IASF), it offers a quantitative and more biologically informative assessment of brain connectivity (Bergamino et al., 2022; Gabusi et al., 2022; Schiavi et al., 2022; Schiavi, Ocampo-Pineda, et al., 2020; Schiavi, Petracca, et al., 2020).

To investigate age-related differences in structural connectivity among various canonical or domain-specific networks, graph theory provides a powerful analytical tool (Fornito et al., 2016; Zhang et al., 2022). Graph theoretical analysis permits the computation of networks at different levels of organization (Fornito et al., 2016; Yeh et al., 2021), using measures classified as (i) local (quantifying properties of individual nodes), (ii) mesoscale (describing interconnected clusters of nodes); and (iii) global (describing whole-brain connectivity properties) (Fornito et al., 2016; Rubinov & Sporns, 2010). At the global scale, graph measures reveal how the brain’s structural wiring facilitates information communication between distant regions and cognitive systems. While structurally connected regions can communicate directly, signal propagation between unconnected nodes requires a sequence of one or more intermediate connections (Zhang et al., 2022). Thus, investigating these measures across and between predefined cognitive systems during development can shed light on the structural mechanisms behind functional expression (Seguin et al., 2019).

In this study, we construct microstructure-informed connectomes and study age-related patterns of local and global structural brain network properties in a typically developing sample aged 8-18 years.

## 2. Materials and methods

### 2.1. Participants

We enrolled a sample of typically developing children and adolescents aged 8-18 years recruited as part of the Cardiff University Brain Research Imaging Centre (CUBRIC) Kids study, with ethical approval from the School of Psychology ethics committee at Cardiff University. Participants and their parents/guardians were recruited via public outreach events, and written informed consent was obtained from the primary caregiver of each child participating in the study. Adolescents aged 16-18 years additionally provided written consent. Children were excluded from the study if they had non-removable metal implants, or a reported history of a major head injury or epilepsy. All procedures were conducted in accordance with the Declaration of Helsinki. A total of 88 children (Mean age = 12.6, SD = 2.9 years) were included in the current study (46 female).

### 2.2. MRI acquisition

Images were acquired on a 3T Siemens Connectom system with ultra-strong (300 mT/m) gradients. As described in (Genc et al., 2020), the protocol comprised: (a) a 3D Magnetization Prepared Rapid Gradient Echo (MPRAGE) for structural segmentation (TE/TR = 2/2300ms; voxel size 1×1×1mm^3^); (b) multi-shell dMRI acquisition (TE/TR = 59/3000 ms; voxel size = 2×2×2mm^3^) with b∈[500, 1200, 2400, 4000, 6000] s/mm^2^ in 30, 30, 60, 60, 60 directions respectively and additional 14 b = 0 s/mm^2^ volumes. Diffusion MRI data were acquired in an anterior-posterior phase-encoding direction, with one additional posterior-anterior volume.

### 2.3. MRI processing

A summary of image processing steps is illustrated in Figure 1. T_1_-weighted data were processed using FreeSurfer version 6.0 (http://surfer.nmr.mgh.harvard.edu) to derive a white matter mask and parcellate the cortical grey matter according to the Destrieux atlas (Destrieux et al., 2010). Next, we registered the Yeo functional atlas (Yeo et al., 2011) in MNI space to each individual subject’s space using a non-linear transformation as implemented in FNIRT of FSL (Smith et al., 2004). This allowed us to obtain eight functionally relevant cortical canonical networks (herein referred to as “Yeo7”) for further interrogation (visual, somatomotor, dorsal attention, ventral attention, limbic, frontoparietal, default mode network, subcortical). Subsequently, we grouped regions of interest (ROIs) from the Destrieux atlas into the eight Yeo atlas networks. To merge the two atlases within each subject, we employed a data-driven approach (see Baum et al. (2017)). Briefly, each parcellated brain region was assigned to one of eight canonical functional brain networks (Yeo et al., 2011) by considering the maximum number of voxels in the intersection between the masks. We ensured that the same overlap was confirmed in the homologous ROIs and for at least 80% of the enrolled subjects, discarding any Destrieux ROIs that did not meet these criteria. The final subdivision can be seen in Figure 2 and Table S2. Finally, we linearly-registered the T_1_-weighted images and the corresponding parcellations on dMRI data using FLIRT (Jenkinson et al., 2002) with boundary-based optimization (Greve & Fischl, 2009).

**Figure 1:**
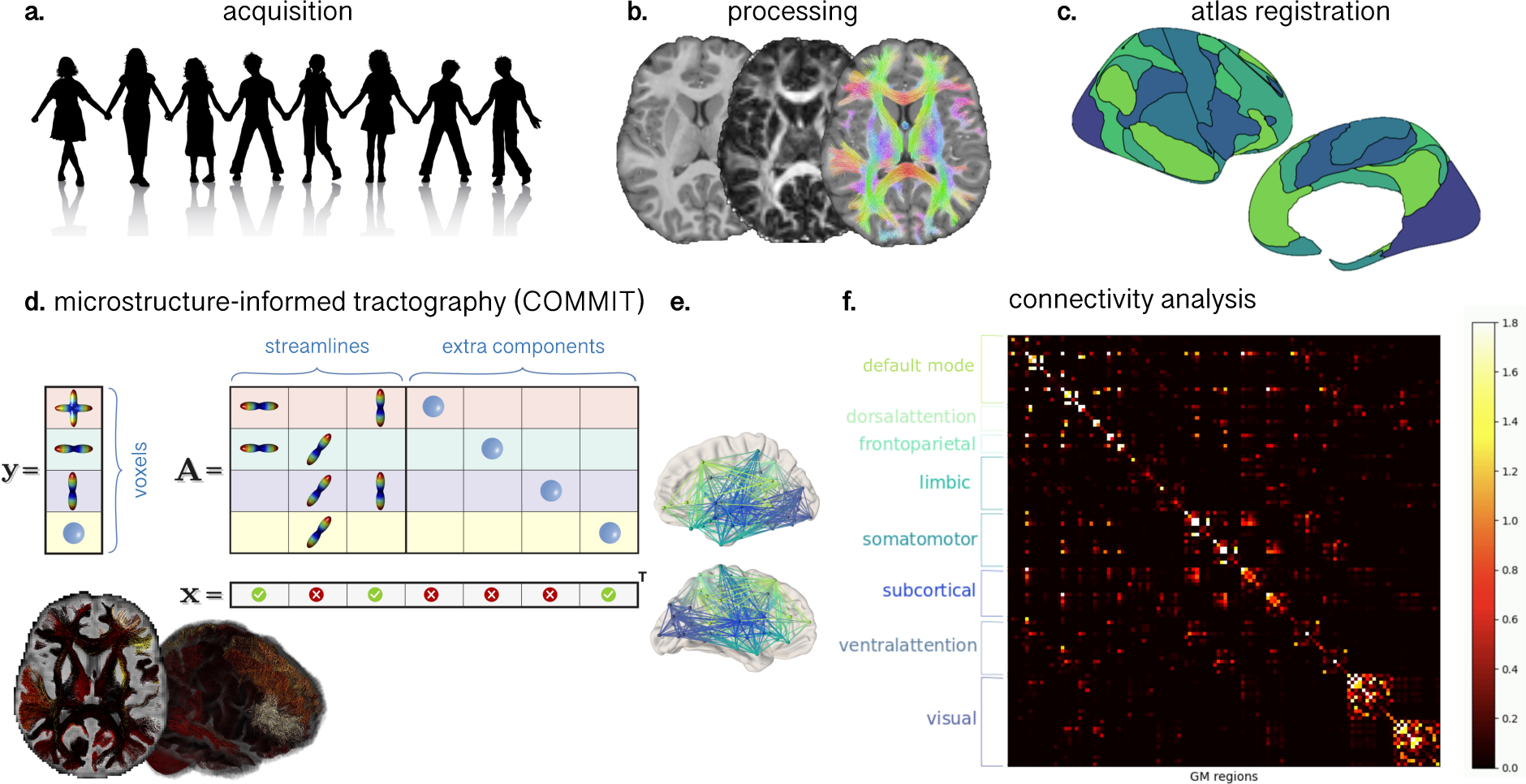
Workflow for constructing structural connectivity networks based on COMMIT derived streamline weights: a) MRI data were acquired in 88 children and adolescents aged 8-18 years; b) T1 and dMRI data were pre-processed; c) canonical cortical networks derived from a functional atlas (Yeo et al., 2011) were co-registered to individual subject space; d) COMMIT (Daducci et al., 2015, 2013) was applied using a stick-zeppelin-ball model to filter out implausible connections, where computed weights reflect the intra-axonal signal fraction of each connection (brighter values = higher IASF); e) interconnected nodes coloured by canonical cortical network; f) connectivity matrix demonstrating connection strength between nodes within in each network (brighter values = higher IASF).

**Figure 2:**
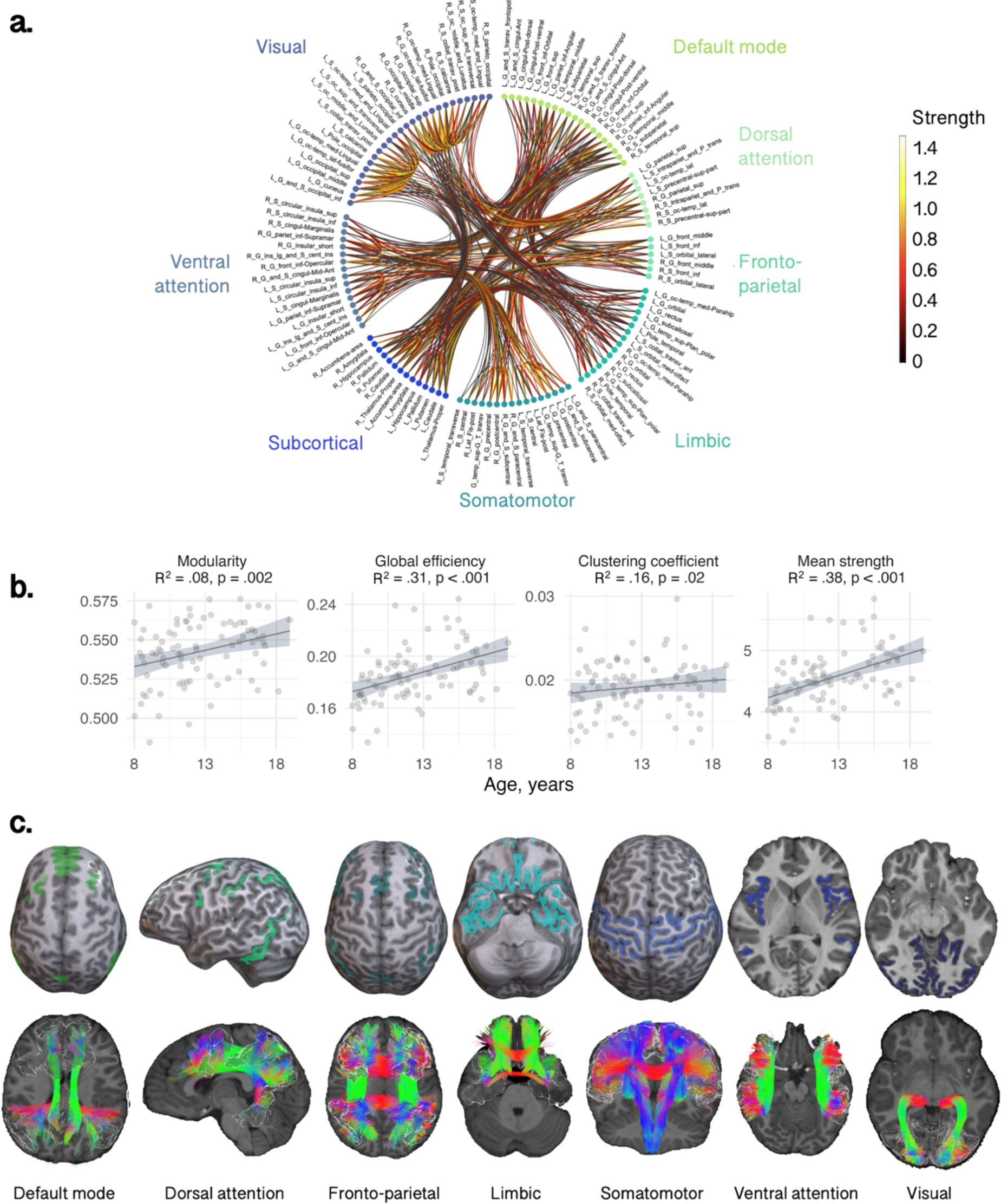
Relationship between age and global network measures computed for the whole connectome realized with Destrieux parcellation. a) The circle plot indicates the connection strength between and within distinct networks obtained using the intra-axonal signal fraction estimated with COMMIT; b) Association between age and network characteristics between networks (R^2^ and p-value); c) Depiction of atlas-derived cortical functional networks and representative white matter tracts traversing these networks, for an 8-year-old female participant.

Diffusion MRI data were pre-processed as detailed in Genc et al. (2020). Briefly the preprocessing pipeline involved FSL (Smith et al., 2004), MRtrix3 (Tournier et al., 2019), and ANTs (Avants et al., 2011) tools using the following steps: denoising (Veraart et al., 2016); slice-wise outlier detection (Sairanen et al., 2018); and correction for drift (Vos et al., 2017); motion, eddy, and susceptibility-induced distortions (Andersson et al., 2003; Andersson & Sotiropoulos, 2016); Gibbs ringing artefact (Kellner et al., 2016); bias field (Tustison et al., 2010); and gradient non-uniformities (Glasser et al., 2013; Rudrapatna et al., 2021). We performed multi-shell multi-tissue constrained spherical deconvolution (MSMT-CSD; Jeurissen et al. (2014)) and generated a whole-brain probabilistic tractogram seeding from the white matter comprising 3 million streamlines (Tournier et al., 2010).

We then applied COMMIT (Daducci et al., 2015, 2013) using a stick-zeppelin-ball model (Panagiotaki et al., 2012) to effectively filter out implausible connections while obtaining the intra-axonal signal fraction for each streamline, as described in Schiavi, Petracca, et al. (2020). For a set of fixed intra- and extra-axonal diffusivities, we assume that the IASF is constant along the streamline. To set the diffusivity parameters in COMMIT, we performed voxel-wise estimations in one younger participant (8-year-old female) and one older participant (17-year-old female). In the white matter, diffusivities had minimal variation between the younger and older participant (Table S1). As a result, for all subjects we set the following diffusivities d_par_=d_par_zep_=1.7×10^−3^ mm^2^/s, d_perp_=0.61×10^−3^ mm^2^/s, d_iso_ in [1.7,3.0]x10^−3^ mm^2^/s for all participants.

For each subject, the connectomes were built using nodes from the individual T1-based Destrieux parcellation by assigning the total IASF associated to each bundle as edge-weights as in Schiavi, Petracca, et al. (2020) and Gabusi et al. (2022). Briefly, for each subject, the microstructure-informed connectomes (i.e., obtained using COMMIT weights reflecting IASF associated to each streamline as entries) were built using the GM parcellation described above and computing the weighted average intra-axonal signal contribution of each bundle:

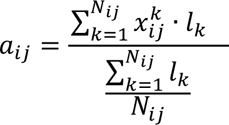

where *i*, *j* are the indices of ROIs connected by the bundle, *N_ij_* is bundle’s number of streamlines, 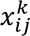 is the weight of the streamline, *k*, obtained by COMMIT, and *l_k_*, its length. In this way, each entry contained the total IASF associated to the bundle given by the weighted average of the streamline contribution multiplied by its length and divided by the average length of the bundle.

### 2.4. Network analysis

To investigate the relationship between network characteristics and age, we used the Brain Connectivity Toolbox for Python (Rubinov & Sporns, 2010) to compute the following network measures:

- modularity (reflecting network segregation);
- global efficiency (corresponding to the average inverse shortest weighted-path length and inversely related to the characteristic path length);
- clustering coefficient (reflecting the degree to which the nodes tend to cluster together); and
- mean strength (corresponding to the average of all the nodal strengths, where the nodal strength is the sum of the weights of links connected to the node).

We computed these global network measures for the entire connectome, as well as within each subnetwork identified within the Yeo7 atlas.

### 2.5. Age-relationships

To investigate age-related patterns of network characteristics across the Yeo7 networks, we applied linear mixed effects modelling using lme4 (Bates et al., 2015) in R (RStudio v3.4.3). We built a linear model which included age (linear term), sex and Yeo7 network as predictors, with intracranial volume (ICV) included as a covariate. We examined four network characteristics (modularity, global efficiency, clustering coefficient, mean strength) and compared the fit of the standard linear model with alternative models that incorporated interaction terms. To identify the most appropriate model, we used the Akaike Information Criterion (AIC) (Akaike, 1974), selecting the model with the lowest AIC as the most parsimonious. Individual general linear models were run to determine age-related differences in specific network characteristics in all eight Yeo7 networks. Evidence for an association was deemed statistically significant when p < .005 (Benjamin et al., 2018).

### 2.6. Feature importance

To identify locally important nodes that contribute to developmental patterns within networks (identified in section 2.5), we performed age-prediction using linear regression and ElasticNet regularization in scikit-learn (i.e., L1 and L2 penalties). We investigated feature importance using the ROIs comprised in each network for age-prediction of local efficiency. First, we randomly split the data into training and validation sets using an 80-20 ratio, resulting in 80% of the data being allocated for training purposes and the remaining 20% for model evaluation (total N=88: 70 training; 18 testing). Then, we performed feature scaling to ensure that all variables were on a similar scale. To assess the generalization performance of the ElasticNet model and to prevent overfitting, we employed a 5-fold cross-validation approach. We performed a grid search to determine the optimal values for the L1 ratio ([0.1, 0.5, 0.7, 0.9, 0.95, 0.99, 1]) based on the regression coefficient (R^2^).

The performance of the model was assessed using the validation dataset. Finally, the features with the largest weight coefficients were extracted to identify specific cortical regions driving age-relationships in local network efficiency.

## 3. Results

### 3.1. Global network characteristics

Linear models revealed a positive relationship between age and modularity (R^2^ = .08, p = .002), global efficiency (R^2^ = 0.31, p < 0.001) and mean strength (R^2^ = .38, p < .001) (Figure 2b). The relationship between age and clustering coefficient was not statistically significant (R^2^ = .16, p = .02). As shown in the circle plot in Figure 2a, we also noted strong intra-regional connectivity and strength within the visual and somatomotor networks, indicating robust interactions among regions within these networks.

To test if specific networks were driving these developmental patterns of network properties, we tested age-by-network interactions using a linear mixed effects model. The various models tested, and the model selection results are summarised in Table S3. The best fitting model for all four graph measures included an age by network by sex interaction term. We observed significant age-by-network interactions in modularity (F = 6.6, p < .001), global efficiency (F = 6.7, p < .001), clustering coefficient (F = 3.3, p = .002), and mean strength (F = 23.9, p < .001). As these results indicated that there were age-related differences in network properties <u>between</u> the networks, we performed subsequent analyses to test for age associations <u>within</u> networks, to discern whether developmental patterns differed regionally. The various networks tested and their corresponding anatomical tractography depictions are illustrated in Figure 2c.

### 3.2. Sub-network characteristics

We identified regional differences in the age-related development of specific sub-networks (Table 1 and Figure 3). Through linear regression analyses within individual networks, we found statistically significant relationships between age and global efficiency in the default mode (R^2^ = .38, p = .001), somatomotor (R^2^ = .28, p < .001) and visual networks (R^2^ = .43, p < .001). Clustering coefficient was positively associated with age in the visual network (R^2^ = .37, p < .001). Moreover, age exhibited a positive association with mean strength in the somatomotor network (R^2^ = .33, p < .001) and the visual network (R^2^ = .46, p < .001). We also observed a negative association between age and modularity in the ventral attention network (R^2^ = .13, p < .001). Overall, our results highlight the distinct age-related developmental patterns in the visual and somatomotor networks.

**Figure 3:**
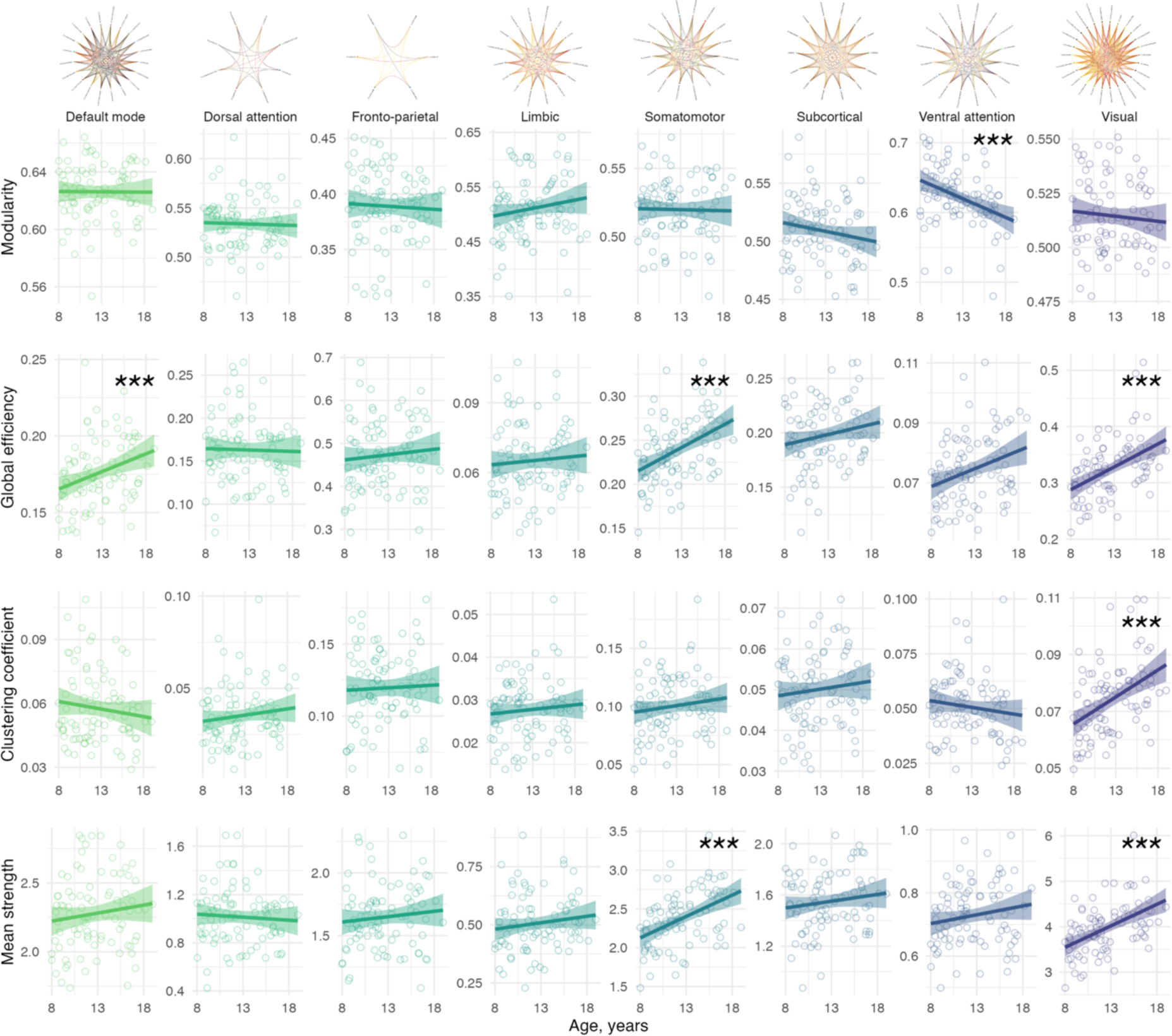
Association between age and network properties within sub-networks. Significant age relationships are annotated (***: p<.005). Top panel represents circle plots of within-network nodes, with brighter yellow connections indicative of higher mean strength.

**Table 1:** Summary statistics for the relationship between age and global sub-network characteristics.

| Network | Modularity |  | Global efficiency |  | Clustering coefficient |  | Mean strength |  |
| --- | --- | --- | --- | --- | --- | --- | --- | --- |
| | $R^2$ | p-value | $R^2$ | p-value | $R^2$ | p-value | $R^2$ | p-value |
| Default mode | .04 | .55 | .38 | **.001** | .10 | .59† | .43 | .13† |
| Dorsal attention | -.03 | .81 | .06 | .41† | .09 | .20 | .06 | .23† |
| Fronto-parietal | .07 | .66 | .03 | .58 | -.01 | .96 | .07 | .51† |
| Limbic | .07 | .14 | .19 | .92 | .14 | .81 | .21 | .53† |
| Somatomotor | .01 | .75 | .28 | **< .001** | .30 | .20 | .33 | **< .001** |
| Subcortical | .08 | .27 | .03 | .26 | .01 | .72 | .02 | .47† |
| Ventral attention | .13 | **< .001** | .19 | .006 | .11 | .47† | .22 | .12† |
| Visual | .11 | .17 | .43 | **< .001** | .37 | **< .001** | .46 | **< .001** |

Note: Adjusted $R^2$ determined using a linear model including age, sex and total intracranial volume. Bold values indicate $p < .005$ . † denotes a significant difference in the slope of the age relationship compared with the visual network.

To confirm that the age-dependence of visual network properties were significantly different from other networks, we performed linear mixed-effects modelling to discern whether age-by-network interactions were significantly different <u>between</u> the visual network and the seven remaining sub-networks. Where the age-relationship in the visual network was significantly stronger than each subsequent network, this is summarised in Table S4 and annotated in Table 1. In summary, the most marked observations were in network strength, where the visual network had a significantly stronger age-dependency compared to each individual network, apart from the somatomotor network which also had a positive relationship with age.

### 3.3. Feature importance of local efficiency

Age prediction of local efficiency in the visual network yielded a regression coefficient of 0.45 (RMSE: 2.2, p=.001, Figure 4a) on the validation set (optimal value for L1=0.1). Feature importance in the visual network identified specific nodes (Figure 4) driving age-related increases in local efficiency. The 10 most sensitive nodes were balanced between hemispheres (5 nodes in right hemisphere, and 5 in the left) and accounted for 75% of variation in total weights (of a total of 26 nodes). Figure 4b summarises the regions ranked by weight, and Figure 4c depicts these regions in axial, sagittal and coronal views in 3D. Nodes with high feature importance for age clustered together, including nodes which form the dorsal (left superior occipital gyrus and middle occipital gyrus and sulcus) and the ventral (right medial occipito-temporal sulcus and gyrus, and right lingual gyrus) visual pathways.

**Figure 4:**
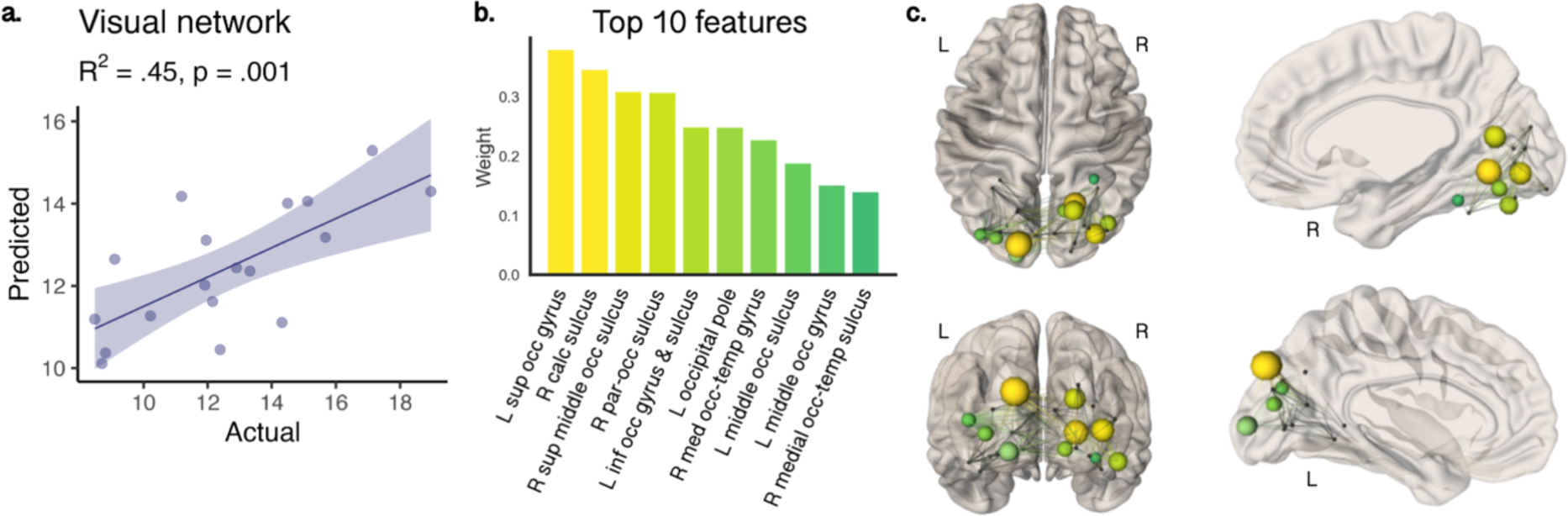
Feature importance for age-prediction of local network efficiency in the visual cortex. A) predicted age was significantly associated with actual age; B) top 10 ranking regions that contributed most to age-related patterns displayed on C) axial, sagittal, and coronal glass brain views, where nodes are scaled and color-coded by weight. Nodes with high feature importance included left superior and middle occipital gyrus and right medial occipito-temporal gyrus.

Age prediction for local efficiency of the somatomotor network yielded a weaker regression coefficient of 0.10 which was not statistically significant (p=.10). Feature importance identified specific regions driving age-related increases in local efficiency. Six nodes balanced between hemispheres (3 nodes in right hemisphere, and 3 in the left) accounted for 70% of the variation in total weights (of a total of 16 nodes). Nodes with high feature importance for age included the bilateral precentral gyrus, right postcentral gyrus, bilateral central sulcus, and left transverse temporal gyrus.

## 4. Discussion

We used microstructure-informed tractography to investigate global and local network characteristics in canonical cortical networks among a group of typically developing children and adolescents. Our study revealed three main findings:

First, whole-brain network-based measures of modularity, global efficiency and mean strength increased with age. This indicates that as children move through adolescence, the shortest path between nodes (in this case, regions from the Destrieux parcellation) decreases, resulting in a more efficient transfer of information. As a result, the nodes tend to cluster together to form hubs, and the strength of each connection increases with age. These findings align with known age-related increases in global efficiency during adolescent development (Baker et al., 2015; Khundrakpam et al., 2013; Koenis et al., 2018; Van den Heuvel & Sporns, 2013). Additionally, previous white matter studies have shown substantial increases in intra-axonal signal fraction with age (Chang et al., 2015; Genc et al., 2020; Palmer et al., 2022), aligning with our observations of age-related increases in mean strength.

Second, sub-network analyses revealed specific networks with substantial age-related differences occurring from childhood to adolescence. In the default mode, somatomotor, and visual networks, global efficiency was higher with older age. Additionally, clustering coefficient was higher with age in the visual network, and mean strength was higher with age in the somatomotor and visual networks. Notably, brain structures, such as the primary visual and somatomotor cortex have highly organized and specialized structures that are closely related to their function, such as discriminating visual features (Wandell, 1999) and performing specific motor functions (Gordon et al., 2023). Together, our findings of age-related maturation of network efficiency and strength suggests a high degree of integration and communication within motor and visual processing regions, potentially reflecting the ongoing maturation of visual information processing and motor coordination capabilities during development. Our specific findings in the visual network align with previously observed temporal patterns of white matter microstructural maturation in the visual cortex (Colby et al., 2011; Genc et al., 2017) which are likely to be closely linked to age-related increases in axon density in humans (Genc et al., 2020) and rodents (Juraska & Willing, 2017).

Age-prediction in the visual cortex pointed to a smaller cluster of five regions per hemisphere that contributed to >75% of the observed age-related differences in local network efficiency. Our data driven approach suggests that nodes in the left dorsal (middle and superior occipital) visual pathway and the right ventral (middle occipito-temporal) visual pathway are driving developmental improvements in local network efficiency. The visual system undergoes early establishment during prenatal development and continues to mature through life (Gogtay et al., 2004; Knudsen, 2004). While myelination in the visual cortex is largely completed by the first year of life (Deoni et al., 2015), recent research indicates that myelination follows a protracted course in ventral temporal cortices (Natu et al., 2019). Ongoing intra-cortical myelination of the ventral temporal cortex may underlie MRI-derived estimates of cortical thinning, previously attributed to synaptic pruning (Gomez et al., 2017; Natu et al., 2019).

The maturation of association visual cortices supports higher level visual processing (e.g. recognising and discriminating objects, motion perception etc.) (Gomez et al., 2018). Our findings align with task-based fMRI studies involving object and shape recognition tasks, which demonstrate protracted development of dorsal and ventral visual pathways (Freud et al., 2019; Ward et al., 2023). These developmental improvements in shape-processing mechanisms likely contribute to microstructure-specific strengthening of global network efficiency and connection strength within the visual network through child and adolescent brain development. The age-related increases in local network efficiency in lateral temporo-occipital cortices may facilitate improvements in visual processing and function in these association cortices.

The myelination of these visual pathways may help to refine and optimize the neural connections and improve visual processing capabilities. Whilst we did not directly study myelination here, the intra-axonal signal fraction explains a significant proportion of the age-related variance in network efficiency and connection strength. Taken together, our findings suggest that the visual cortex undergoes protracted development through childhood and adolescence. While our study primarily focuses on white matter microstructure for exploring graph-based measures, our observations of higher efficiency and connection strength with older age is predominantly due to ongoing microstructural maturation in the visual cortex.

### 4.1. Methodological advantages of the current approach

We employed a data-driven approach to establish correspondence between a structural parcellation and functional atlas in each participant (Baum et al., 2017). This involved selecting the maximum number of voxels in the intersection between a smaller cortical region with its corresponding larger functional network. By ensuring that this overlap was consistent with the homologous ROIs and in at least 80% of the participants, we generated canonical cortical networks for the basis of regional graph-based analyses.

One of the significant advantages of the COMMIT framework is its ability to assign specific microstructural properties to individual tractography-reconstructed streamlines, which sets it apart from conventional (voxel-wise or vertex-wise) approaches where complex intra-voxel heterogeneity can bias estimates (Schilling et al., 2022). By allowing a distribution of microstructural values to be assigned to a voxel, i.e., the number of values is equal to the number of unique streamlines passing through the voxel and retained for analysis, COMMIT offers a more complete estimation of microstructural properties. In the context of graph theory, we are better equipped to capture the dynamic strengthening and weakening of connections with maturation over childhood and adolescence. Overall, the COMMIT framework offers a more nuanced and detailed characterization of microstructural properties along individual streamlines, countering complex intra-voxel heterogeneity, making it a powerful tool for a more meaningful assessment of brain connectivity (Gabusi et al., 2022; Schiavi et al., 2022; Schiavi, Ocampo-Pineda, et al., 2020; Schiavi, Petracca, et al., 2020).

### 4.2. Limitations and future directions

It is important to acknowledge that certain functional networks utilised in our study here contain fewer nodes than others, potentially influencing our interpretations. Although we adopted a robust method to generate reproducible cortical nodes for each functional network, it resulted in some networks having a small number of nodes.

While there is a certain relationship between brain structure and function, structure-function coupling occurs in a spatially-dependent hierarchical manner (Baum et al., 2020). The brain is a complex and dynamic organ, with function influenced by a variety of factors, including structural organisation (Chamberland et al., 2017) and neural activity. Whilst the aforementioned factors may help explain why we did not observe an age dependence of network-based measures of brain connectivity in regions known to remodel in adolescence (e.g. the fronto-parietal network), it is known that functional networks that are in close range demonstrate stronger white matter connectivity (Hermundstad et al., 2013), which may explain why our findings of global efficiency and mean strength were confined to the somatomotor and visual networks. On the note of the fronto-parietal network, despite running a ‘gold-standard’ dMRI pre-processing pipeline, susceptibility-induced distortion artefacts may introduce an additional source of variance into the diffusion MRI data, especially in fronto-parietal regions with an air/bone interface such as the nasal cavity.

Future work characterising the developing connectome using biologically meaningful mathematical models of brain connections are promising (Akarca et al., 2023; Seguin et al., 2023). Combining task-based or resting-state fMRI with microstructure-informed connectomes may better elucidate structure-function coupling across the developing brain (Suárez et al., 2020). Recent updates to the COMMIT framework offer the opportunity to incorporate additional imaging contrasts, such as myelin-sensitive contrasts, leading to improved delineation of anatomically accurate whole-brain tractography (Leppert et al., 2023; Schiavi et al., 2022).

## 5. Conclusion

Incorporating microstructural information into network analyses has shed light on distinct regional age-related development of brain networks. Notably, we observed unique characteristics within the visual network throughout development, supporting its ongoing maturation, reaffirming previously reported patterns of protracted development in the dorsal and ventral visual pathways. Overall, our study demonstrates the power of microstructure-informed tractography to decipher intricate developmental patterns, reinforcing the potential for deepening our understanding of brain connectivity and development.

## 6. Supporting information

### Acknowledgments

We are grateful to the participants and their families for their participation in this study. We thank Umesh Rudrapatna and John Evans for their support with image acquisition protocols, Isobel Ward for assistance with data collection, Joseph Yang for scientific discussions, and Greg Parker for contributions to data pre-processing and model fitting pipelines. Image credit (Figure 1a) by kjpargeter on Freepik.

### Funding

The data were acquired at the UK National Facility for In Vivo MR Imaging of Human Tissue Microstructure funded by the EPSRC (grant EP/M029778/1), and The Wolfson Foundation. SS received funding from the University of Verona Internationalisation Programme 2019 (Action 4C). This research was funded in whole, or in part, by a Wellcome Trust Investigator Award (096646/Z/11/Z) and a Wellcome Trust Strategic Award (104943/Z/14/Z). CMWT was supported by a Veni grant (17331) from the Dutch Research Council (NWO) and the Wellcome Trust [215944/Z/19/Z]. For the purpose of open access, the author has applied a CC BY public copyright licence to any Author Accepted Manuscript version arising from this submission.

### Authors’ Contributions

S.S., S.G. and D.K.J. conceptualized the problem. S.S., S.G., M.C. and C.T. analyzed the MRI data. S.G. and E.R. acquired all MRI data. S.G and M.C. performed statistical analyses. A.D. and D.J., supervised and raised funding for this project. S.S., S.G. and D.K.J. wrote the original draft of the manuscript. S.S., S.G., M.C., C.T., E.R., A.D., and D.K.J. reviewed and edited the manuscript.

### Code and data availability

The code for COMMIT is open source and freely available at https://github.com/daducci/COMMIT.

### Disclosures

Declarations of interest: SG, MC, CT, ER, AD, DKJ declare no conflict of interest. SS is employed by ASG Superconductors S.p.A. but there is no financial interest related to this work.

## 8. Supplementary

### 8.1. Tables

**Table S1:** Diffusivity parameters estimated in a white matter mask for one younger (8-year-old) and one older (17-year-old) participant. Values are reported as mean (SD)

| | $d_a$ | $d_{par}$ | $d_{perp}$ |
| --- | --- | --- | --- |
| Younger | 2.27 (0.71) | 2.01 (0.57) | 0.61 (0.28) |
| Older | 2.35 (0.62) | 1.71 (0.58) | 0.62 (0.27) |

**Table S2:**
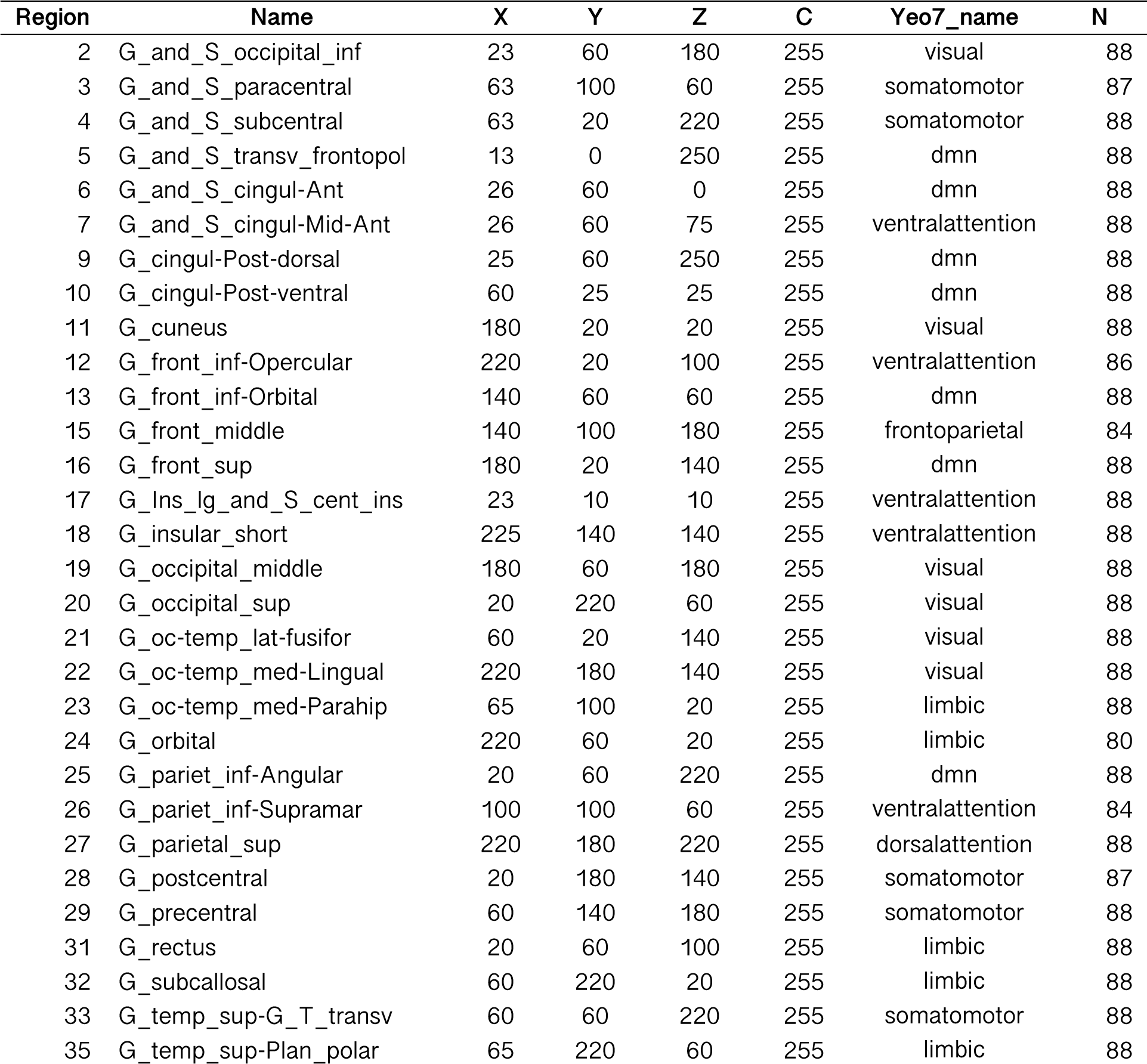
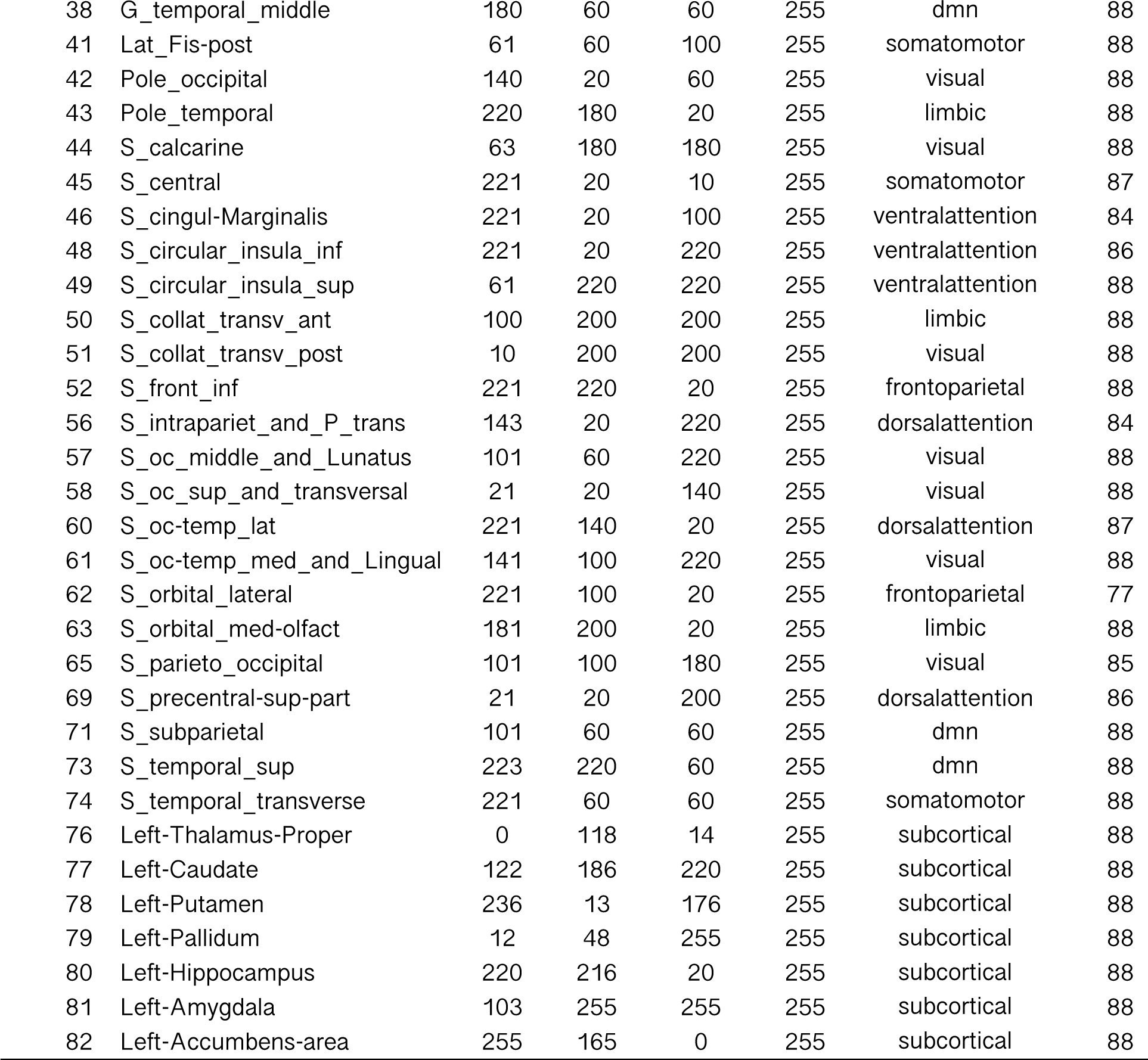
Regions from the Destrieux parcellation assigned to each canonical cortical network. Results for left hemisphere shown (equivalent in right hemisphere). Only nodes overlapping the same network in >80% of participants were included in the analysis.

**Table S3:** Results of mixed-effect model selection for first level global graph network analysis. Values reported are Akaike Information Criterion (AIC) of each model fit.

| Model | Modularity | Global Efficiency | Clustering Coefficient | Mean Strength |
| --- | --- | --- | --- | --- |
| M1a | -2816.78 | -2549.70 | -3792.68 | 244.49 |
| M2a | -2821.03 | -2553.69 | -3796.30 | 237.85 |
| M3a | -2832.02 | -2565.74 | -3795.91 | 167.79 |
| M4a | -2860.55 | -2575.29 | -3814.42 | 90.46 |
| M1b | -2825.11 | -2569.99 | -3801.41 | 215.66 |
| M2b | -2826.42 | -2570.17 | -3802.19 | 214.00 |
| M3b | -2840.35 | -2586.03 | -3804.65 | 138.97 |
| M4b | **-2865.94\*** | **-2591.77\*** | **-3820.31\*** | **66.60\*** |

Note: Bold indicates lowest AIC for each graph measure; \* indicates if the age by network term was significant at $p < .005$
Footnote: Models tested are as follows:
M1a <- lmer(measure ~ age + sex + network + (1|ID), REML=FALSE, data=data)
M2a <- lmer(measure ~ age \* sex + network + (1|ID), REML=FALSE, data=data)
M3a <- lmer(measure ~ age \* network + sex + (1|ID), REML=FALSE, data=data)
M4a <- lmer(measure ~ age \* sex \* network + (1|ID), REML=FALSE, data=data)
M1b <- lmer(measure ~ age + sex + network + ICV + (1|ID), REML=FALSE, data=data)
M2b <- lmer(measure ~ age \* sex + network + ICV + (1|ID), REML=FALSE, data=data)
M3b <- lmer(measure ~ age \* network + sex + ICV + (1|ID), REML=FALSE, data=data)
M4b <- lmer(measure ~ age \* sex \* network + ICV + (1|ID), REML=FALSE, data=data)

**Table S4:** Results from comparison of age-associations of graph measures with reference to the visual network. Bold values indicate networks which have significantly different slopes to the age-relationship in the visual network, generated using linear mixed efforts models.

| Network | Global efficiency |  | Clustering coefficient |  | Mean strength |  |
| --- | --- | --- | --- | --- | --- | --- |
|  | t | p-value | t | p-value | t | p-value |
| *Visual (reference)* |  |  |  |  |  |  |
| Default mode | -1.65 | .10 | -2.91 | **.004** | -4.08 | **5E-05** |
| Dorsal attention | -2.96 | **.003** | -1.17 | .24 | -5.25 | **2E-07** |
| Fronto-parietal | -1.64 | .10 | -1.66 | .10 | -3.91 | **1E-04** |
| Limbic | -2.04 | .04 | -1.87 | .06 | -4.01 | **7E-05** |
| Somatomotor | -0.60 | .55 | -1.16 | .25 | -1.78 | .08 |
| Subcortical | -2.19 | .03 | -2.19 | .03 | -4.55 | **6E-06** |
| Ventral attention | -1.93 | .05 | -3.05 | **.002** | -4.34 | **2E-05** |
| *Somatomotor (reference)* |  |  |  |  |  |  |
| Default mode | -1.05 | .29 | -1.05 | .29 | -2.30 | .02 |
| Dorsal attention | -2.36 | .02 | -2.36 | .02 | -3.48 | **< .001** |
| Fronto-parietal | -1.04 | .30 | -1.04 | .30 | -2.13 | .03 |
| Limbic | -1.45 | .15 | -1.45 | .15 | -2.23 | .03 |
| Subcortical | -1.60 | .11 | -1.60 | .11 | -2.78 | .006 |
| Ventral attention | -1.33 | .18 | -1.33 | .18 | -2.56 | .011 |
| Visual | 0.60 | .55 | 0.60 | .55 | 1.78 | .08 |

Note: model used was the best fitting model deduced from Table S3.

